# Drug synergy scoring using minimal dose response matrices

**DOI:** 10.1101/2020.10.30.362103

**Authors:** Petri Mäkelä, Si Min Zhang, Sean G Rudd

**Affiliations:** Department of Oncology-Pathology, Science For Life Laboratory, Karolinska Institutet, Stockholm, Sweden

**Keywords:** Cancer, Combination therapy, Precision medicine, Synergy, Antagonism, Dose-response matrix, Dose-response landscape, Checkerboard assay

## Abstract

**Objective:** Combinations of pharmacological agents are essential for disease control and prevention, offering many advantages over monotherapies, with one of these being drug synergy. The state-of-the-art method to profile drug synergy in preclinical research is by using dose-response matrices in disease-appropriate models, however this approach is frequently labour intensive and cost-ineffective, particularly when performed in a medium- to high-throughput fashion. Thus, in this study, we set out to optimise a parameter of this methodology, determining the minimal matrix size that can be used to robustly detect and quantify synergy between two drugs.

**Results:** We used a drug matrix reduction workflow that allowed the identification of a minimal drug matrix capable of robustly detecting and quantifying drug synergy. These minimal matrices utilise substantially less reagents and data processing power than their typically used larger counterparts. Focusing on the antileukemic efficacy of the chemotherapy combination of cytarabine and inhibitors of ribonucleotide reductase, we could show that detection and quantification of drug synergy by three common synergy models was well-tolerated despite reducing matrix size from 8×8 to 4×4. Overall, the optimisation of drug synergy scoring as presented here could inform future medium- to high-throughput drug synergy screening strategies in pre-clinical research.

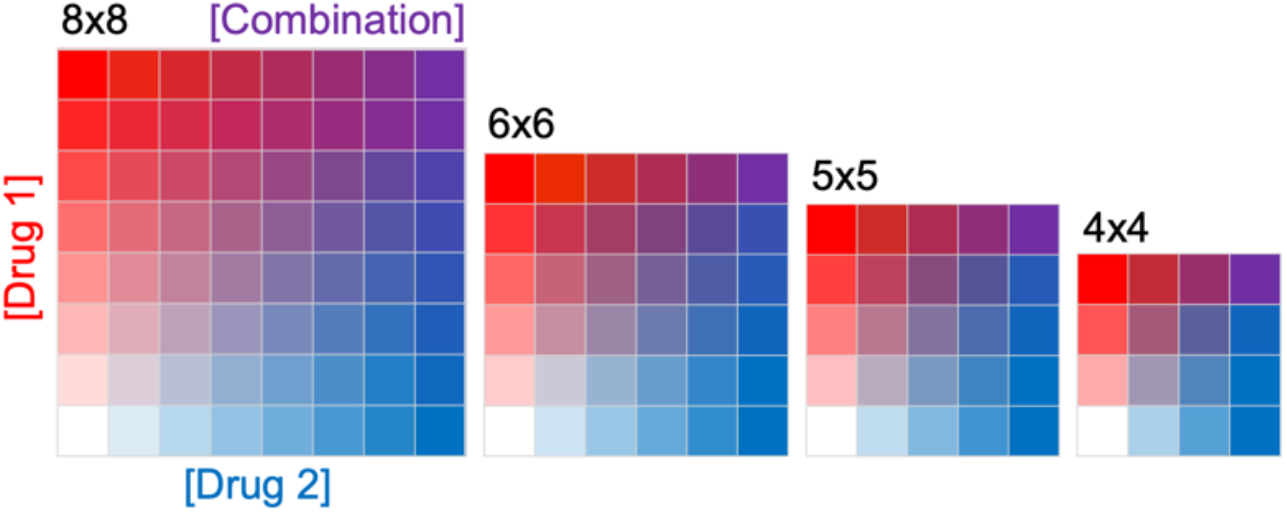

## Introduction

Current treatment regimens for many different diseases utilise combinations of pharmacological agents, and this is especially true in the treatment of cancer. The rationale behind the use of two or more drugs in cancer therapy is to enhance cancer cell killing, reduce treatment toxicity, and prevent the onset of treatment resistance. There is ample clinical evidence documenting the benefit of this approach for cancer patients (1), with one of the first being in acute leukaemias (2). As oncology continues to move towards personalised treatment strategies, be it with traditional cytotoxic chemotherapies or with targeted therapies, ultimately these agents will be used in a combination regimen, and it is important to ensure these combinations are developed in a rational manner (3). It is thus critical to robustly assess drug-drug interactions at the pre-clinical stage and to translate this knowledge into the clinic.

One parameter of combination therapy that is routinely the focus of pre-clinical research is drug synergy/antagonism scoring (4). Although there is a lack of nomenclature standardisation (5,6), synergy can be broadly defined as a combination effect that is stronger than expected from the sum of the drugs individual effects, whilst antagonism is a combination effect that is less active than the additive effect. Although drug synergy is not necessarily required for clinical benefit (7), with an additive effect being sufficient to cure in some instances (8), synergy/antagonism scoring remains an important parameter to evaluate when designing combination therapies or working to understand the mechanisms underpinning current treatment regimens.

The most straight-forward and cost-effective setting in which to assess drug-drug interactions is in cultured cancer cell lines, and the information generated here can be translated into more complex cancer models. There are a number of methodologies to assess drug-drug interactions in cancer cell lines, ranging from those requiring minimal effort but yielding little information, to those which can be more labour intensive but generate a comprehensive profile of drug-drug interaction (4,5,9). These methods range from (i) testing of single drug doses alone and in combination, (ii) the use of dose gradients in which drug combinations are tested at a single fixed ratio, and (iii), the use of dose-response matrices (also referred to as a checkerboard assay) which provide complete dose-response information for the tested combination. The latter approach provides the most comprehensive profile of drug-drug interaction, but requires more datapoints, and thus reagents, to achieve this, which can limit throughput potential. Thus, in this study, we set out to optimise drug synergy scoring using dose-response matrices by questioning at which point reducing the matrix size would compromise on robust drug synergy scoring.

## Materials and methods

### Cell lines

The THP-1 cell line used in this study is a CRISPR/Cas-9 control clone, the generation of which has been described previously (10). Cells were cultured in IMDM medium (#12440053, Gibco), supplemented with 10% FCS (#10500064, Gibco) and penicillin-streptomycin (#15070063, Gibco) at 37°C and 5% CO_2_ in a humidified incubator. Cells were routinely monitored and tested negative for mycoplasma using MycoAlert (#LT07-318, Lonza).

### Compounds

Ara-C (#C1768, Sigma-Aldrich) and dF-dC (#G6423, Sigma-Aldrich) were prepared at 10 mM stock concentration in DMSO (#23486, VWR Chemicals) and stored at −20°C. HU (#H8627, Sigma-Aldrich) was prepared fresh at 50 mM stock concentration in DMSO.

### Drug combination assay

The proliferation inhibition and drug synergy assay has been described previously (11). Compound dispensing in flat, clear-bottomed 384-well microplates (#3764, Corning) and DMSO volume normalisation was performed using the D300e Digital Dispenser (Tecan) with the aid of the Synergy Wizard in the D300e Control Software. Plate layouts included two columns of DMSO to be used as positive (cells suspension supplemented with DMSO) and negative controls (media only with DMSO). Cell suspensions (20 000 cells/ml) were dispensed into these plates using a MultiDrop (Thermo Fisher Scientific), dispensing 50 μl per well (thus 1000 cells per well). Plates were then placed in a pre-warmed humidity chamber consisting of a plastic box containing damp paper towels and incubated for 4 days at 37°C and 5% CO_2_ in a conventional humidified incubator. To quantify remaining viable cells, 10 μl resazurin solution (#R17017, Sigma-Aldrich; prepared to 0.06 mg/ml in PBS) was added to each well and further incubated for 6 hours prior to fluorescence measurements (530/590 nm, ex/em) using a Hidex Sense Microplate Reader. Fluorescent intensity of each well was normalised to the average of the control wells on the same plate to calculate relative cell viability values. For synergy analysis, relative cell viability measurements from duplicate wells were averaged and analysed using the web-based tool SynergyFinder (12,13). Synergy summary scores were derived from the average of the synergy scores across the entire dose-response landscape. Data visualisation and statistical testing was performed using Prism 8 (GraphPad Software).

## Results

Reducing the size of a drug matrix vastly reduces the wells used in a microwell plate (**Table 1**), but it remains unclear which matrix size can robustly detect and quantify drug synergy. As an example of a synergistic interaction between two anti-cancer drugs by which to address this question, we chose the deoxycytidine analogue cytarabine (ara-C) and the ribonucleotide reductase (RNR) inhibitors hydroxyurea (HU) or gemcitabine (dF-dC), the latter of which is also a deoxycytidine analogue. Synergistic killing of cancer cells by this drug combination has been documented for decades (reviewed in (11)), and thus, we utilised this example in the following study to investigate which size dose-response matrix can still robustly detect and quantify this drug-drug interaction.

**Table 1.** Comparison of matrix sizes.

| Matrix | Wells used in a 384-wp<sup>a</sup> | Matrices per 384-wp (incl. controls<sup>b</sup>) |
| --- | --- | --- |
| 8x8 | 64 | 5 |
| 6x6 | 36 | 9 |
| 5x5 | 25 | 14 |
| 4x4 | 16 | 22 |

<sup>a</sup> Abbreviations: 384-wp, 384-well microplate.
<sup>b</sup> Typical experimental setup includes one column (or equivalent) each of positive and negative controls, and thus this is accounted for in the matrices per plate calculation.

In this workflow, outlined in **Fig. 1**, we began by determining the concentration range required to produce a complete dose-response curve for each drug in each cell line by performing monotherapy dose-response analyses. Having a complete monotherapy dose-response is ideal for comprehensively profiling drug-drug interaction when compounds are then evaluated in combination. However, in some instances this may not be possible due to the activity range of the compound or compound solubility, which may limit the maximum concentration that can be tested. After selecting the concentration ranges to be evaluated, we then designed an experiment in which several drug matrix sizes were tested on the same microtiter plate in duplicate. Matrix sizes began at 8×8 and was reduced to 6×6, 5×5, and 4×4, each having the same highest and lowest compound concentration with doses between equally, logarithmically spaced. Each matrix included a dose-response of each drug alone, together with no compound (i.e. solvent only), thus an 8×8 matrix includes 7 doses each tested in combination (49 combinations in total) whilst a 4×4 includes 3 doses each tested in combination (9 combinations in total). The acute myeloid leukaemia (AML) cell line, THP-1, was then seeded upon these differing dose-response matrices and, following a 4-day incubation, resazurin reduction used to measure the remaining metabolically viable cells. Relative cell viabilities were then calculated and analysed via the SynergyFinder web-application (12,13) using 3 alternate drug-drug interaction models, zero interaction potency (ZIP) (14), bliss independence (15), and highest single agent (HSA) (16). This experiment was repeated four times on different days and the data subsequently combined, shown in **Fig. 2**.

**Figure 1.**
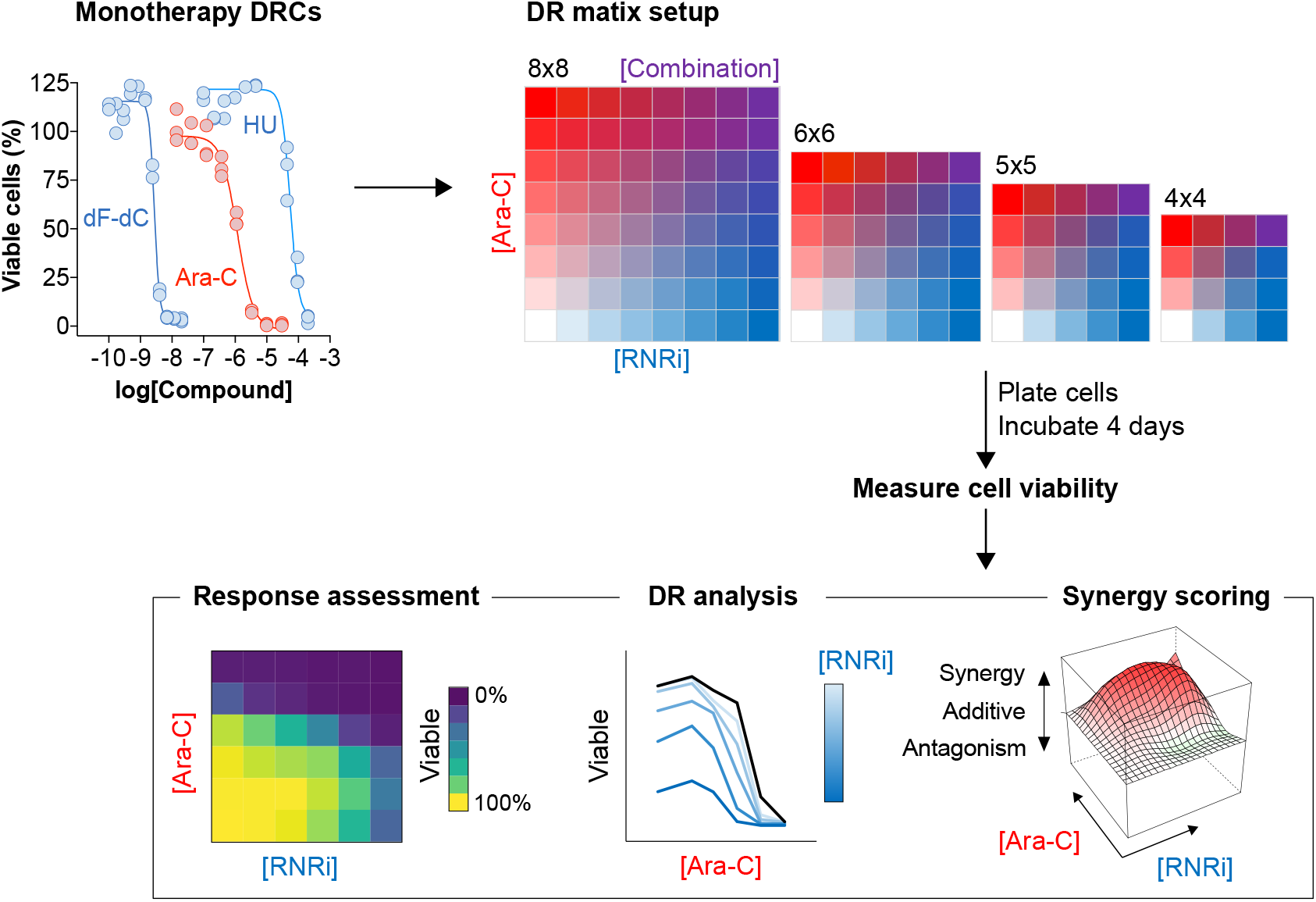
Overview of experiment to evaluate minimal dose-response matrices. Schematic detailing experiment conducted in this report. Chemotherapeutics cytarabine (ara-C) and ribonucleotide reductase inhibitors (RNRi) hydroxyurea (HU) or gemcitabine (dF-dC) are first evaluated in monotherapy dose-response curve (DRC) analyses before being combined in different dose-response (DR) matrix sizes. Following incubation with cells, response to drug treatment is assessed before DR analysis and synergy scoring.

**Figure 2.**
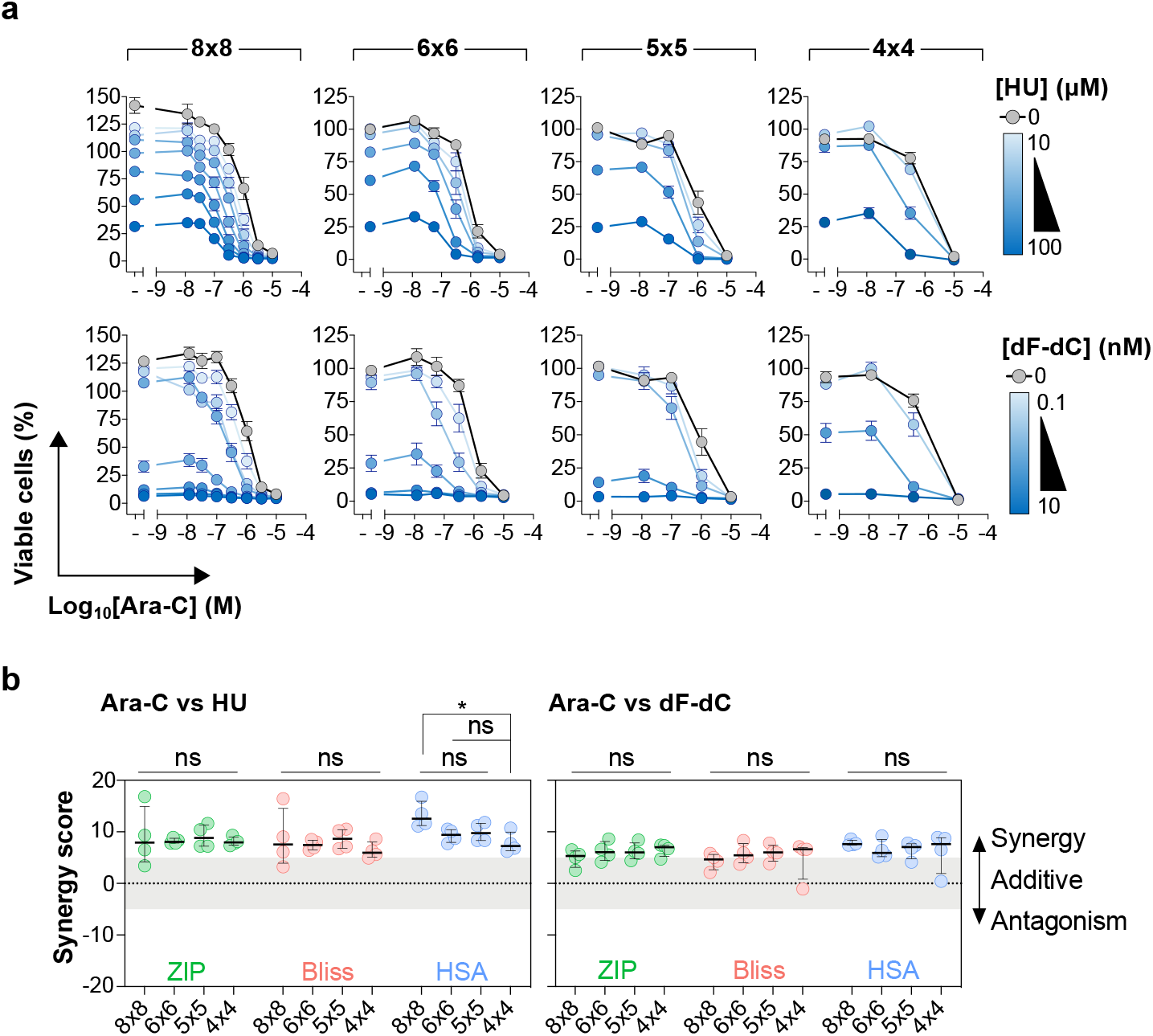
Dose-response and synergy scores produced from different matrix sizes. **A.** Relative cell viability plotted as a function of ara-C concentration at differing hydroxyurea (HU) or gemcitabine (dF-dC) doses. Mean values from four independent experiments plotted, error bars indicate s.e.m. **B.** Drug synergy plots for ara-C and the indicated RNR inhibitor, either HU or dF-dC, using the different synergy models. Each data point indicates an average synergy score from a single doseresponse matrix experiment performed in duplicate. Zero, >0 or <0 corresponds to additive, synergy or antagonism, respectively, whilst >5 indicates strong synergy. The horizontal line and the error bars indicate the median and interquartile range, respectively, from four independent experiments. Statistical testing was carried out using the non-parametric Kruskal-Wallis test: ns, not significant; *, p > 0.05.

We first plotted relative cell viability as a function of ara-C concentration with increasing RNRi dosage (**Fig. 2a**). Regardless of matrix size, the dose-dependent sensitisation of THP-1 cells to ara-C by either HU or dF-dC could be clearly observed. However, although the ara-C sensitisation was visible in all matrix sizes, the resolution of the dose-response data was obviously reduced in the smaller matrices. We next compared the synergy summary scores from the different matrix sizes (**Fig. 2b**). We observed that all matrices tested could detect a synergistic interaction between ara-C and HU or dF-dC in THP-1 cells. Comparing the synergy values within each synergy model using the non-parametric Kruskal-Wallis test, we found that the vast majority of matrix sizes showed no significant difference in the quantity of synergy measured. Altogether, 36 comparisons were made and only 1 gave a statistically significant difference, which was the 8×8 matrix compared with the 4×4 using the HSA model (p=0.025), but this significant difference was not observed using the alternate synergy models.

## Discussion

In this study, we set out to scale down the size of dose-response matrices used to assess drug synergy, as although this method produces the most comprehensive dataset, it is often cost-prohibitive. Comparison of 8×8, 6×6, 5×5, and 4×4 matrices revealed no consistent difference in detecting and quantifying synergy between two chemotherapeutic agents. Thus, the minimal 4×4 and 5×5 matrices were capable of quantifying drug synergy to an extent equal to the larger matrices, despite requiring substantially less wells in a microtiter plate (**Table 1**). Accordingly, this would reduce the running cost of this approach considerably, and allow more combinations or cell models to be screened on the same microtiter plate, which could be an important consideration for medium- to high-throughput drug combination screens.

In support of the utility of minimal drug matrices, we recently used this approach in testing drug combinations in a panel of AML cell lines to identify a biomarker for drug synergy, which we confirmed in ex vivo experiments in patient-derived AML blasts (11). Furthermore, a pseudo-5×5 matrix (monotherapy dose-responses performed separately to a 4×4 combination matrix) has been successfully deployed in a large-scale drug combination screen in cancer cell lines (17), and the NCI-ALMANAC study also contains 4×4 drug matrices (18,19).

Several alternate approaches have been suggested with the aim to reduce the cost of high-throughput drug combination screening, such as using a cross-combination design (20) or utilising a sub-matrix design coupled with machine learning, which is readily accessible through a web-based application (21). The approach suggested in this study is not mutually exclusive with those previously reported, and perhaps future studies could evaluate the use of the cross-combination or sub-matrix design based upon a minimal dose-response matrix to potentially further increase throughput of drug combination screens.

## Limitations

A principle limitation of this study is that it utilises only three chemotherapeutics combined into two combinations which are tested in one cancer cell model, by which to optimise the methodology, and of course, there are infinitely more pharmaceutical agents and combinations that can be assessed. Thus, it is possible that conclusions made here may not be translated to other combinations or preclinical cancer models; this remains to be tested. However, the workflow outlined in this study could be first utilised with the drug combinations and/or disease models of interest in order to inform further experiments.

Regarding the minimal matrices, whilst the 4×4 matrix could robustly detect and quantify synergy to the same extent as larger matrices, resolution of dose-response information was reduced, which could be an important consideration when setting up a drug combination experiment. This is especially true given that some synergy metrics (such as ZIP (14)) requires accurate curve fitting to the datapoints (although this was not a limitation in the 4×4 matrices shown in this study). Furthermore, the approach of using a minimal matrix requires pre-screening of each compound as a monotherapy in order to determine the concentration range to be tested in the dose-response matrix, which may not always be possible depending upon the drug combination screening setup. Another consideration is that this method utilises automation and liquid handling equipment to increase technical accuracy and this equipment may not be readily available due to cost, however the technical robustness provided by an automated setup is a significant advantage. Given the reduction of dose-response resolution by the minimal 4×4 matrix, a compromise could be to run drug combination screens with a 5×5 matrix, as this provides a good balance between (i) reagents consumed, (ii) robust detection and quantification of synergy, and (iii), dose-response resolution.

## Declarations

### Availability of data and materials

data and materials are available from the corresponding author upon reasonable request

### Author contributions

S.G.R. conceptualised and supervised the study. S.G.R and P.M. designed experiments, P.M. performed experiments, and P.M. and S.G.R. analysed the data. S.M.Z. and S.G.R. prepared figures and wrote the manuscript.

## Acknowledgements

We thank Ingrid Almlöf and Elisée Wiita for advice on microtiter plate-based assays and Cynthia Paulin for discussion of data.

## References

1. Frei E, Eder JP. Principles of Dose, Schedule, and Combination Therapy. In: Kufe DW, Pollock RE, Weichselbaum RR, et al., editors. Holland-Frei Cancer Medicine. 6th edition. Hamilton (ON): BC Decker; 2003. Chapter 44. Available from: https://www.ncbi.nlm.nih.gov/books/NBK12635/

2. Frei E, Holland JF, Schneiderman MA, Pinkel D, Selkirk G, Freireich EJ, et al. A comparative study of two regimens of combination chemotherapy in acute leukemia. Blood. 1958;13(12):1126–48.

3. Boshuizen J, Peeper DS. Rational Cancer Treatment Combinations: An Urgent Clinical Need. Mol Cell; 2020;78(6):1002–18.

4. Pemovska T, Bigenzahn JW, Superti-Furga G. Recent advances in combinatorial drug screening and synergy scoring. Curr Opin Pharmacol. 2018;42:102–10.

5. Foucquier J, Guedj M. Analysis of drug combinations: current methodological landscape. Pharmacol Res Perspect. 2015;3(3):e00149–11.

6. Meyer CT, Wooten DJ, Lopez CF, Quaranta V. Charting the Fragmented Landscape of Drug Synergy. Trends in Pharmacological Sciences. 2020; 41(4):266–280.

7. Palmer AC, Sorger PK. Combination Cancer Therapy Can Confer Benefit via Patient-to-Patient Variability without Drug Additivity or Synergy. Cell. 2017;171(7):1678–1682.

8. Palmer AC, Chidley C, Sorger PK. A curative combination cancer therapy achieves high fractional cell killing through low cross-resistance and drug additivity. eLife. 2019;8:679–36.

9. Vlot AHC, Aniceto N, Menden MP, Ulrich-Merzenich G, Bender A. Applying synergy metrics to combination screening data: agreements, disagreements and pitfalls. Drug Discovery Today. 2019;24(12):2286–98.

10. Herold N, Rudd SG, Ljungblad L, Sanjiv K, Myrberg IH, Paulin CBJ, et al. Targeting SAMHD1 with the Vpx protein to improve cytarabine therapy for hematological malignancies. Nat Med. 2017;23(2):256–63.

11. Rudd SG, Tsesmetzis N, Sanjiv K, Paulin CB, Sandhow L, Kutzner J, et al. Ribonucleotide reductase inhibitors suppress SAMHD1 ara-CTPase activity enhancing cytarabine efficacy. EMBO Molecular Medicine. 2020; 12(3):e10419.

12. Ianevski A, He L, Aittokallio T, Tang J. SynergyFinder: a web application for analyzing drug combination dose-response matrix data. Bioinformatics. 2017;33(15):2413–5.

13. Ianevski A, Giri AK, Aittokallio T. SynergyFinder 2.0: visual analytics of multi-drug combination synergies. Nucleic Acids Res. 2020;48(W1):W488–93.

14. Yadav B, Wennerberg K, Aittokallio T, Tang J. Searching for Drug Synergy in Complex Dose–Response Landscapes Using an Interaction Potency Model. CSBJ. 2015;13(C):504–13.

15. Bliss CI. The Toxicity Of Poisons Applied Jointly. Annals of Applied Biology. 1939;26(3):585–615.

16. Gaddum JH. Pharmacology. 1940.

17. O'Neil J, Benita Y, Feldman I, Chenard M, Roberts B, Liu Y, et al. An Unbiased Oncology Compound Screen to Identify Novel Combination Strategies. Molecular Cancer Therapeutics. 2016;15(6):1155–62.

18. Holbeck SL, Camalier R, Crowell JA, Govindharajulu JP, Hollingshead M, Anderson LW, et al. The National Cancer Institute ALMANAC: A Comprehensive Screening Resource for the Detection of Anticancer Drug Pairs with Enhanced Therapeutic Activity. Cancer Res. 2017;77(13):3564–76.

19. Close DA, Wang AX, Kochanek SJ, Shun T, Eiseman JL, Johnston PA. Implementation of the NCI-60 Human Tumor Cell Line Panel to Screen 2260 Cancer Drug Combinations to Generate >3 Million Data Points Used to Populate a Large Matrix of Anti-Neoplastic Agent Combinations (ALMANAC) Database. SLAS Discov. 2019;24(3):242–63.

20. Malyutina A, Majumder MM, Wang W, Pessia A, Heckman CA, Tang J. Drug combination sensitivity scoring facilitates the discovery of synergistic and efficacious drug combinations in cancer. PLoS Comput Biol. 2019;15(5):e1006752.

21. Ianevski A, Giri AK, Gautam P, Kononov A, Potdar S, Saarela J, et al. Prediction of drug combination effects with a minimal set of experiments. Nat Mach Intell. 2019;1(12):568–77.

